# Variation of carbon, nitrogen and phosphorus content in fungi reflects their ecology and phylogeny

**DOI:** 10.1101/2024.01.31.578150

**Authors:** Matěj Pánek, Tereza Vlková, Tereza Michalová, Jan Borovička, Leho Tedersoo, Bartosz Adamczyk, Petr Baldrian, Rubén Lopéz-Mondéjar

**Affiliations:** Laboratory of Environmental Microbiology, Institute of Microbiology of the Czech Academy of Sciences, Prague, Czech Republic; Institute of Geology of the Czech Academy of Sciences, Prague, Czech Republic; Nuclear Physiscs Institute of the Czech Academy of Sciences, Husinec-Řež, Czech Republic; Mycology and Microbiology Center, University of Tartu, Tartu, Estonia; Natural Resources Institute Finland, Helsinki, Finland; Department of Soil and Water Conservation CEBAS-CSIC, Campus Universitario de Espinardo, Murcia, Spain

**Keywords:** Fungal biomass composition, Nutrient stoichiometry, Nutrient content variation, Phylogenetic signal, Ecological traits

## Abstract

Fungi are an integral part of the nitrogen and phosphorus cycling in trophic networks, as they participate in biomass decomposition and facilitate plant nutrition through root symbioses. Nutrient content varies considerably between the main fungal habitats, such as soil, plant litter or decomposing dead wood, but there are also large differences within habitats. While some soils are heavily loaded with N, others are limited by N or P. One way in which nutrient availability can be reflected in fungi is their content in biomass. In this study, we determined the C, N, and P content (in dry mass) of sporocarps of 214 fungal species to inspect how phylogeny and membership in ecological guilds (soil saprotrophs, wood saprotrophs, and ectomycorrhizal fungi) affect the nutrient content of fungal biomass. The C content of sporocarps (415 ± 25 mg g^-1^) showed little variation (324-494 mg g^-1^), while the range of N (46 ± 20 mg g^-1^) and P (5.5 ± 3.0 mg g^-1^) contents was within one order of magnitude (8-103 mg g^-1^ and 1.0-18.9 mg g^-1^, respectively). Importantly, the N and P contents were significantly higher in the biomass of soil saprotrophic fungi compared to wood saprotrophic and ectomycorrhizal fungi. While the average C/N ratio in fungal biomass was 11.2, values exceeding 40 were recorded for some fungi living on dead wood, typically characterized by low N content. The N and P content of fungal mycelium also showed a significant phylogenetic signal, with differences in nutrient content being relatively low within species and genera of fungi. A strong correlation was found between N and P content in fungal biomass, while the correlation of N content and the N-containing fungal cell wall biopolymer – chitin showed only weak significance. The content of macronutrients in fungal biomass is influenced by the fungal life style and nutrient availability and is also limited by phylogeny.

## Introduction

Fungi are important decomposers of organic matter in terrestrial trophic networks (Bai et al., 2021; Lindahl et al., 2002). They inhabit environments where sufficient carbon (C) is usually available, but this is not the case for the main macronutrients, nitrogen (N) and phosphorus (P), which are often limiting. While in agricultural soils N and P deficiency is often overcome by fertilization, in most natural ecosystems available N or P sources are limited (Du et al., 2020). While both C and N are primarily present in soil organic matter, P is also mobilised in soil by mineral weathering (Larsen, 1967), while N also comes from wet or dry atmospheric deposition (van Breemen and van Dijk, 1988), often enhanced by human activities such as combustion processes. The access of fungi to N and P in the environment varies depending on their lifestyle (Põlme et al., 2020; Root, 1967), reflecting the C resources used by fungi. The most important ecological guilds of fungi (Põlme et al., 2020) in forest trophic networks are (i) wood-decomposing saprotrophs/parasites that accumulate C from decaying wood, (ii) soil and litter saprotrophs that acquire C through litter decomposition, and (iii) mycorrhizal fungi that acquire C from their plant hosts.

Of the substrates used by fungi, wood from living or dead trees contains the least N and P (Watkinson et al., 2006), whereas soil, and especially topsoil, is richer in these nutrients (Mooshammer et al., 2014). In a typical forest, N or P content may limit fungal activity in dead wood even when content of nutrients in soil is high (Piché-Choquette et al., 2023). Thus, wood saprotrophs (WS) must cope with nutrient limitation in their habitat. Soil saprotrophic (SS) and ectomycorrhizal (EcM) fungi utilize soil N and P, ectomycorrhizal fungi utilize C provided by their host trees in exchange for N (Lindahl et al., 2007; Rayner and Boddy, 1989; Smith and Read, 2008), and therefore their C acquisition costs are reduced. In principle, the benefits of EcM symbiosis appear to be highest in N-limited soils (Johnson, 2010), because the provision of N by fungi to plants leads to higher primary production, thereby increasing C availability to fungi. The ratio between N and P also appears to be very important for mycorrhizal fungi, as higher N supply promotes photosynthetic activity of host plants, which in turn have a higher capacity to exchange C for P (Johnson, 2010).

Increased N availability associated with human activities has been shown to increase the ratio of bacterial and fungal biomass in soil and inhibit microbial respiration and abundance (Ramirez et al., 2012; Zhou et al., 2017). Soil fungal communities are also affected by increased N availability due to anthropogenic deposition (Baldrian et al., 2023), including changes in abundance and balance between nutrient guilds and species within guilds (Baldrian et al., 2022; van der Linde et al., 2018).

Due to both natural variation and anthropogenic deposition, the total amounts of C, N, and P, as well as their ratios, vary substantially among soils (Cleveland and Liptzin, 2007). The amounts and ratios of these elements in fungal biomass also appear to vary among species, both under laboratory conditions and in nature (Brabcová et al., 2018; Camenzind et al., 2021). The analysis of nutrient content in fungal biomass under real conditions is methodically challenging, as it is virtually impossible to separate the hyphae of individual fungal species from their habitat matrix – plant litter, soil, or wood. However, assuming that nutrient content may be reflected in the biomass of fungi growing under laboratory conditions (Camenzind et al., 2021), it is also evident in the biomass that forms the fungal sporocarps (Vogt et al., 1981).

For ectomycorrhizal fungi, it has been suggested that the C:N ratio in their biomass remains relatively constant (Mooshammer et al., 2014), as a result of providing a ‘surplus’ of N to their tree hosts (Kranabetter et al., 2019). In saprotrophic fungi, the storage of P and C is expected in their biomass (Kalač, 2019; Quinché, 1997), while any excess nitrogen, if present, is likely excreted (Mooshammer et al., 2014). Although nutrient content can vary even between different parts of the same individual (Trocha et al., 2016), it can be assumed that differences between ecological guilds should be much more pronounced.

Occasionally, higher concentrations of N and therefore lower C:N ratios in sporocarps have been observed in saprotrophic fungi compared to EcM fungi (Vogt et al., 1981; Zhang and Elser, 2017). This may indicate the lower costs of C in EcM fungi. However, it is well known that fungal species exhibit a preference for environments with specific (high or low) N availability (van der Linde et al., 2018). It should be noted that the nutrient content in fungal biomass is not only a response variable to species traits and environmental conditions; it also influences the fate of microbial biomass in the ecosystem. The rate of decomposition of fungal mycelia in forest soils has been reported to increase with the N content (Brabcová et al., 2018), and it is evident that N is a primary target for guilds of microorganisms responsible for decomposing fungal biomass (López-Mondéjar et al., 2020; Starke et al., 2020). While N is present in many biomolecules, including proteins and nucleic acids (Marzluf, 1996), it also constitutes a significant component of the fungal cell wall polysaccharide chitin (Lenardon et al., 2010), whose content in fungal cell walls varies (Blumenthal and Saul, 1957) and under starvation conditions this compound may undergo enzymatic self-degradation (Gruber and Seidl-Seiboth, 2012; White et al., 2002). However, the chitin content in biomass may to some extent determine the N requirement of each fungal species and potentially correlate with the N content in their biomass.

Substantial differences in the accessibility of C, N, and P by various ecological guilds of fungi raise the question of whether their amounts and ratios are conserved in fungal groups that obtain these nutrients from soil, plant litter, deadwood, or the roots of their host plants. In these environments, the relative content of C, N, and P can vary over more than an order of magnitude (Piché-Choquette et al., 2023). The composition of fungal biomass may also be conserved phylogenetically, at least over short phylogenetic distances (Mouginot et al., 2014), reflecting traits such as the proportion of protein or chitin content in their cell walls. Alternatively, the N and P content in fungal biomass may vary within each species, simply reflecting the accessibility of N and P in each individual site where the fungus is found.

We anticipate that the ecological traits of fungi, particularly those related to the mode and source of C and nutrient acquisition, are the primary drivers of the nutrient content in their biomass. This is because the differences in nutrient limitation between habitats are too significant to be overcome.

Some level of phylogenetic conservation is also likely due to the evolutionary preservation of biochemical traits and the similarity in the ecology of closely related taxa. We analyzed the sporocarps of fungi from temperate forests to determine their C, N, and P contents, aiming to assess the contributions of ecology and phylogeny to biomass composition. We assume that N and P contents are higher in soil saprotrophic fungi than in wood-decomposing and ectomycorrhizal fungi. This is due to the low nutrient concentration in wood in the former case and the exchange of nutrients for carbon with the host plant in the latter case.Additionally, we expect a positive correlation between chitin content and N content in fungal biomass. Furthermore, our study was intended to provide data on the stoichiometric ratios of main macronutrients in the biomass of macrofungi, offering reliable information on this biogeochemically important feature.

### Materials and Methods Samples

Fresh fungal sporocarps were collected in the Czech Republic, Germany, and Estonia between the years 2014 and 2020. Where possible, sporocarps were transferred to liquid nitrogen after collection and kept frozen or immediately transferred to the laboratory and stored at -18 to -20 °C. Subsequently, whole sporocarps were lyphilised and homogenized using a mortar and pestle or a rotary mill to obtain a fine powder, followed by freeze-drying and storage at -18°C until further processing. In total, 282 samples of fungal sporocarps were included in the analysis (Supplementary Table 1).

### Identification of fungal samples and phylogenetic analysis

Fungi were preliminarily identified during collection based on morphology of fungal sporocarps. This preliminary identification was later confirmed or corrected based on the analysis of DNA from the collected specimens. DNA extraction from 250 mg of freeze-dried powdered sporocarp biomass was performed using the DNeasy Qiagen PlantMini kit (Qiagen) in duplicate per specimen. For species identification, the primer combination ITS0F/ITS4 was used to amplify the regions of ITS1, 5.8S, and ITS2 of the rDNA; for phylogeny construction, the primer combination LR0R/LR7, amplifying a part of 28S rDNA, was used (Hibbett and Vilgalys, 1993; Schneider et al., 2015; Schoch et al., 2012; Vilgalys and Hester, 1990; White et al., 1990). The PCR mix contained 200 μM dNTPs (Bioline), 0.5 μM of both LR0R and LR7 primers, or 0.4 μM of both ITS0F and ITS4 primers, 0.02U/μl of Q5 High-Fidelity DNA polymerase (New England Biolabs), 5 ng/μl of DNA in 1 × Q5 Reaction Buffer enriched by 1 × Q5 HighGC Enhancer, and 9.03 μM BSA (GeneOn). The reaction parameters were identical for both DNA regions except for the annealing temperatures. The cycling conditions for ITS were as follows: 98 °C for 30 s; 30 cycles of 98° for 10 s, annealing for 50 °C for 30 s, 72° for 30 s; then 72 °C for 2 min, while for 28S rDNA these conditions were: 95 °C for 5 min.; 35 cycles of 95 ° for 1 min, 52 ° for 1 min 72 ° for 1 min; then 72 °C for 10 min. The PCR product was sequenced using Sanger sequencing in an external facility (SeqMe, Czech Republic).

All DNA sequences were checked for accuracy and manually edited if necessary. They were deposited in GenBank under the accession numbers OR602209-OR602444, OR625684-OR625708, PP102200 and PP102454-PP102678 (Supplementary Table 1). For precise species determination, the sequences of the ITS rDNA regions were compared with the NCBI database using the BLASTn algorithm. For the construction of a phylogenetic tree, the 28S sequences were aligned using the Clustal-W algorithm in BioEdit (Hall, 1999). Phylogenetic trees for the entire dataset and for partial alignments grouping taxonomic or functional groups were constructed using MrBayes (Ronquist et al., 2012) with the standard setup of the evolutionary model. The number of MCMC generations was set individually for particular datasets between 5×10^5^ and 5.5×10^6^ to reach an effective number of independent draws from their posterior distribution higher than 100, and the Potential Scale Reduction Factor (comparing the estimated between-MCMC chain variance with the within-chain variance) close to 1.0.

Fungal species were assigned to taxa at different taxonomic levels and categorized into ecological guilds based on their primary lifestyles, as described in (Põlme et al., 2020). We distinguished wood saprotrophs (WS, including plant pathogens of living trees), soil saprotrophs (SS, encompassing both soil and litter saprotrophs), and ectomycorrhizal species (EcM).

### Analysis of chemical composition

The N and C content in samples were determined at the Institute of Botany of the Czech Academy of Sciences using the automatic element composition analyzer Flash 2000 (Thermo Scientific, USA).

Samples (particle size < 0.1mm, heaped 10-30 mg) were introduced into the burning tube and combusted at 1000 °C in a stream of pure O_2_. Moisture was removed on a separating column, and the content of separated oxides was determined by the conductivity detector. The signal was analyzed using the Eager Xperience software (Thermo Scientific) (Nelson and Sommers, 1996).

The determination of P content in sporocarps was conducted by burning 50 mg of the sample in a porcelain crucible for 6 hours at 550 °C. Subsequently, 1 mL of HNO_3_ was added and heated to a boil. The samples were then transferred to a volumetric flask and replenished with distilled H_2_O to a total volume of 50 mL. The concentration of P was measured colorimetrically (Ohno and Zibilske, 1991). Due to the limited fungal biomass in certain samples resulting from the small size of sporocarps, this analysis was conducted on only a subset of the total sample set.

For selected samples representing various ecological groups of fungi, chitin content was determined using a method based on acid hydrolysis of chitin, producing glucosamine. The concentration of glucosamine was measured after derivatization with FMOC-Cl (9-fluorenylmethyl-chloroformate). The measurement was conducted via HPLC (Arc HPLC, Waters, USA) with Hewlett Packard ODS Hypersil (5 μm, 250 × 4.6 mm) column and detected with a fluorescence detector (Adamczyk et al., 2020).

### Statistical analysis

The statistical analyses were conducted in the R environment (R Core Team, 2022). Descriptive statistics (mean, standard deviation) for the content of C, N, and P were calculated using the R package ‘psych’ (Revelle, 2023), coefficient of variation was calculated as the ratio of standard deviation to mean expressed as a percentage. Comparisons were made considering taxonomic placement and ecological guild membership. The results were visually presented using boxplots and line charts created with the R package ‘ggplot2’ (Wickham, 2016). The significance of differences in the C, N, P, and chitin content between taxa at various taxonomic levels and between ecological guilds was evaluated using the Kruskal-Wallis test, followed by the Dunn test, implemented with the ‘FSA’ R-package (Ogle et al., 2022). The strength of correlations between variables was assessed using Pearson’s r with the ‘ggpubr’ R-package (Kassambara, 2020).

The phylogenetic trees, along with the data on nutrient levels for all samples, were utilized to calculate Pagel’s lambda (λ_P_), expressing the value of phylogenetic signal (0 = closely related samples are not more similar than distant ones, 1 = total determination of the trait by phylogenetic relations between samples, Molina-Venegas and Rodríguez, 2017) for the content of N and the C:N ratio. The calculation was carried out using the ‘phytools’ R-package (Revell, 2012) with the lambda method. To rule out the possibility of the influence of the number of samples (n) in respective categories on the λ_P_ value, a correlation test between λ_P_ and n was performed using Pearson’s r, implemented with the ‘ggpubr’ R-package (Kassambara, 2020).

## Results

The fungal sporocarps samples belonged to 13 orders, predominantly covering *Basidiomycota*, with some species of *Ascomycota*. Among the 282 samples, there were 212 species from 82 fungal genera and 53 families. These were classified according to Põlme et al. (2020) into EcM (154 samples), SS (73 samples), and WS (55 samples; Supplementary Table 1, Figure 1).

### C, N, P, and chitin content in fungal biomass

The C content in fungal sporocarps ranged from 29.8% to 49.4% and showed relatively little variation (41.5 ± 2.6, n = 271). The N content was more variable, spanning more than one order of magnitude from 0.83% to 10.3% (4.6 ± 2.0, n = 271). The highest N content, exceeding 10%, was observed in the soil saprotrophic fungi *Agaricus depauperatus* and *Agaricus phaeolepidotus*, as well as in the ectomycorrhizal species *Porphyrellus porphyrosporus*. The P content ranged from 1.0 mg g^-1^ to 18.9 mg g^-1^ (5.5 ± 3.0, n = 155), with the highest content above 1.8% observed in the soil saprotrophic species *Agaricus campestris* and *Lacrymaria glareosa*. The average ratio of C:N:P across the whole dataset was 99:9:1, but the variation of stoichiometric ratios among samples was wide: C/N ranged from 3.6 to 50.0 (11.2 ± 6.4, n = 271), and C/P ranged from 20.8 to 349 (99 ± 61, n = 145; Supplementary Table 1).

There was a significant negative correlation between the contents of C and P (Pearson’s r = -0.226, p = 0.0063, Figure 2) within whole dataset, while the content of N and P showed a stronger positive correlation (r = 0.478, p < 0.001, Figure 2). However, when considering ecological guilds, this correlation was found to be significant only within the EcM fungi group, which had the highest number of observations (r = 0.285, p = 0.0048). There was a correlation (r = 0.298, p = 0.0239, Figure 2) between chitin and N, as well as between chitin and the C:N ratio (r = -0.290, p = 0.0282) when all samples are considered together.

**Figure 1.**
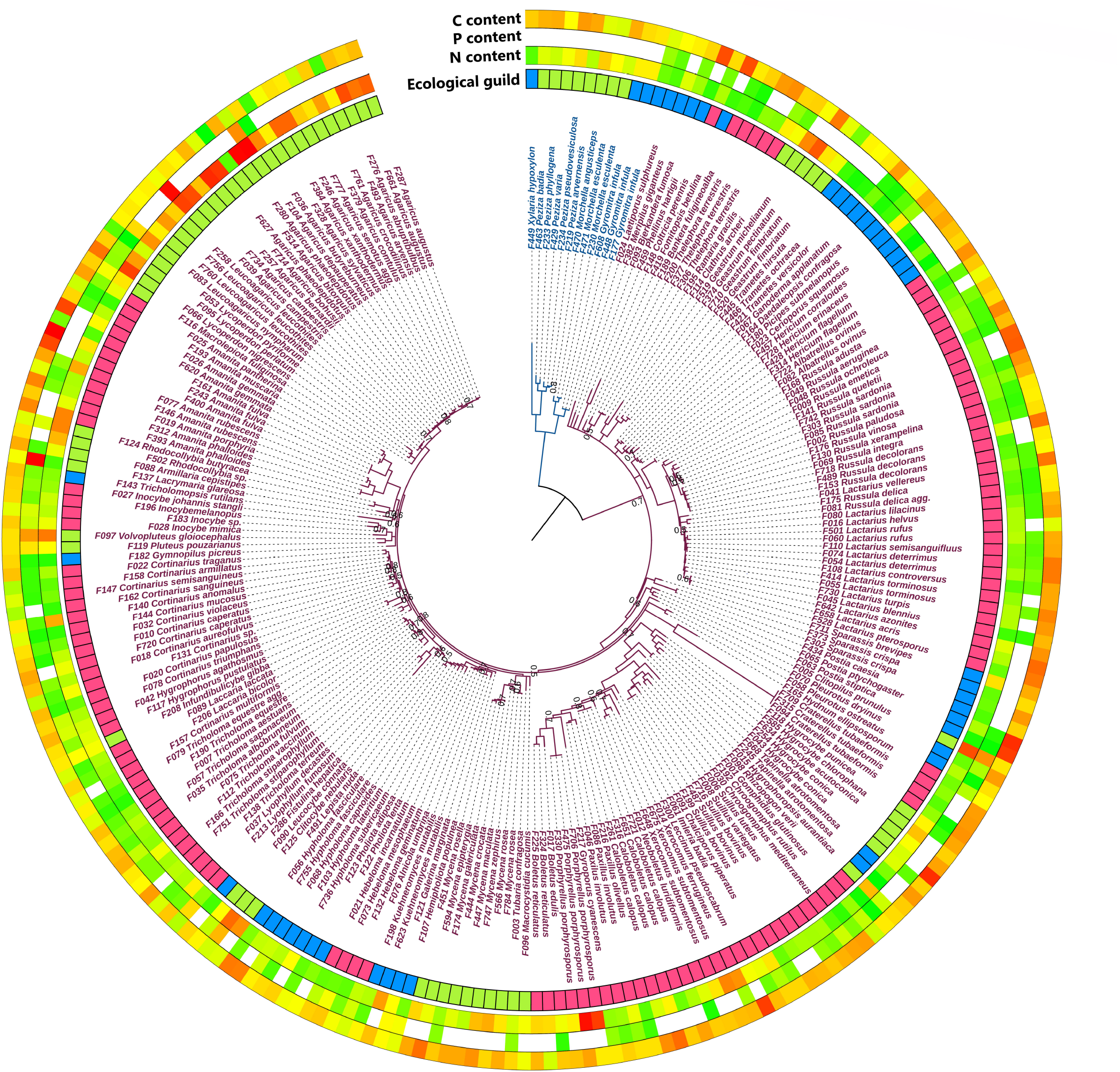
Phylogenetic tree of 282 fungal samples based on the 28S rDNA region. Ecological guild: blue – wood saprotrophs, red – ectomycorrhizal fungi, green – soil saprotrophs. Colors of the outer rings indicate content from low (green) to high (red). Species names of the *Basidiomycota* are purple, and the *Ascomycetes* are blue. At tree nodes, only values of posterior probabilities less than or equal to 0.8 are given.

**Figure 2.**
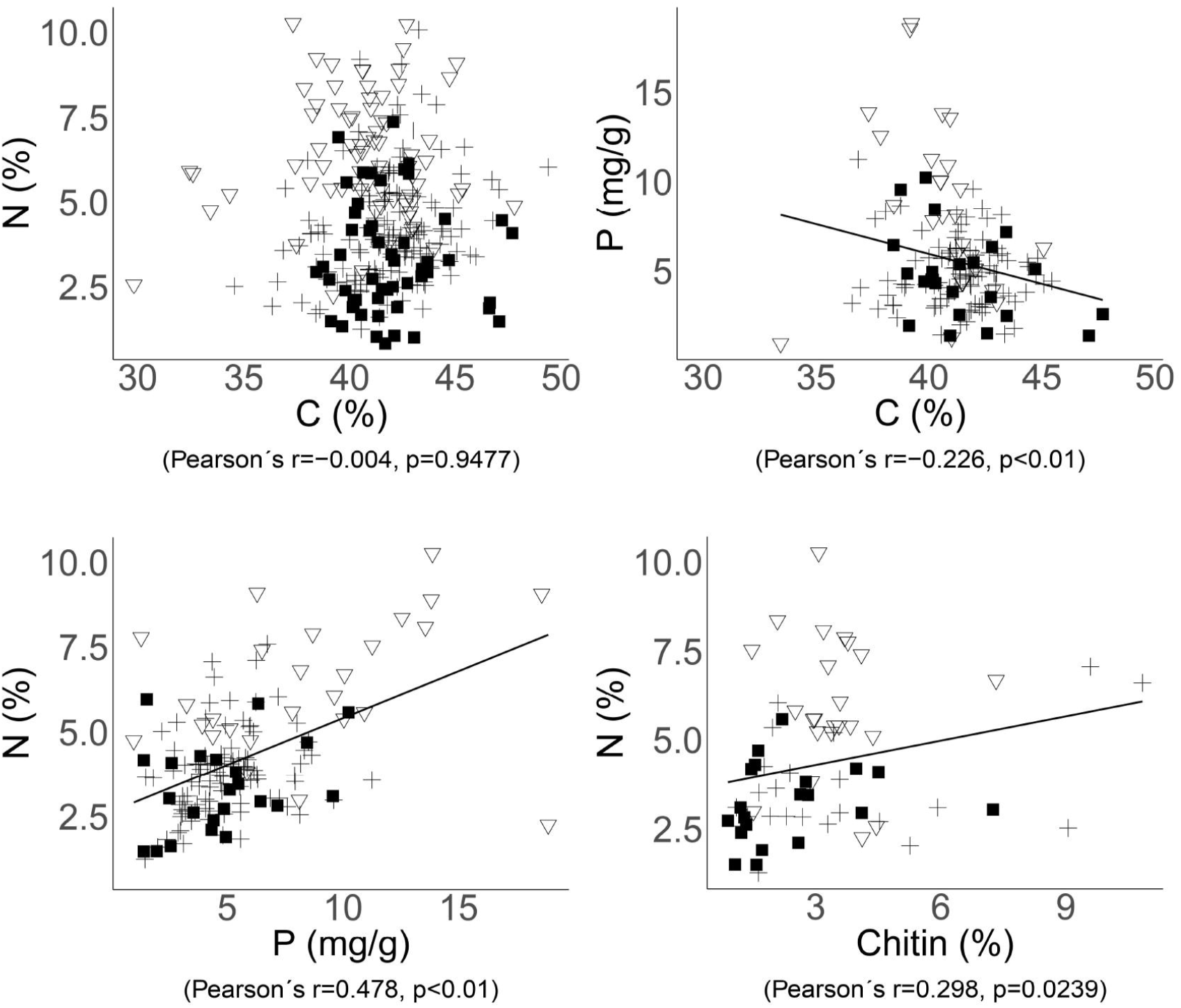
Relationships between the C, N, P, and chitin contents in fungal biomass. Significant trends across the whole dataset are indicated by correlations with Pearson’s correlation coefficients and p-values. Each point represents one species of ectomycorrhizal (cross), soil saprotrophic (triangle), or wood saprotrophic (square) fungi.

The chitin content in fungal sporocarps varied between 0.9% and 10.8% (3.1 ± 2.0, n = 70). The content of chitin was smallest in wood saprotrophs (2.4% ± 1.6% of dry mass), significantly less than in soil saprotrophs (3.4% ± 1.2%). In ectomycorrhizal fungi, chitin content varied more widely (3.6% ± 2.8%; Supplementary Table 2), and this group also harbored species with the highest chitin content around 10% - *Amanita citrina* and *Cortinarius anomalus*. The chitin content showed a significant positive correlation with N content in fungal biomass (Pearson’s r = 0.298, p = 0.0239; Figure 2), despite the fact that N contained in chitin represented just a small proportion of the total N in fungal biomass - between 4.56% and 7.36%.

### Nutrient contents in fungi of contrasting ecology

Interestingly, the nutrient content in fungal biomass differed between ecological groups of fungi (Table 1, Figure 3). Although no significant differences were found for C content between fungal groups (Figure 3), the mean content of N and P in the biomass of soil saprotrophs was significantly higher, by approximately 80%, than in wood saprotrophs (p < 0.05, Kruskal-Wallis test). The content of N in ectomycorrhizal fungi was significantly lower than in soil saprotrophs yet higher than in wood saprotrophs (p < 0.05). Regarding P content, the mean values in ectomycorrhizal and wood saprotrophs fungi were similar, but significantly different in soil saprotrophs. When comparing the ratios between elements, we observed that the C:N ratio exhibited an inverse trend to the N content. Specifically, wood saprotrophs displayed the highest C:N ratio values, whereas soil saprotrophs exhibited the lowest values. Despite the values for C:P and N:P ratios being highly variable between the three ecological guilds, our results showed that the C:P ratio was significantly different between wood saprotrophs and soil saprotrophs.

**Table 1.**
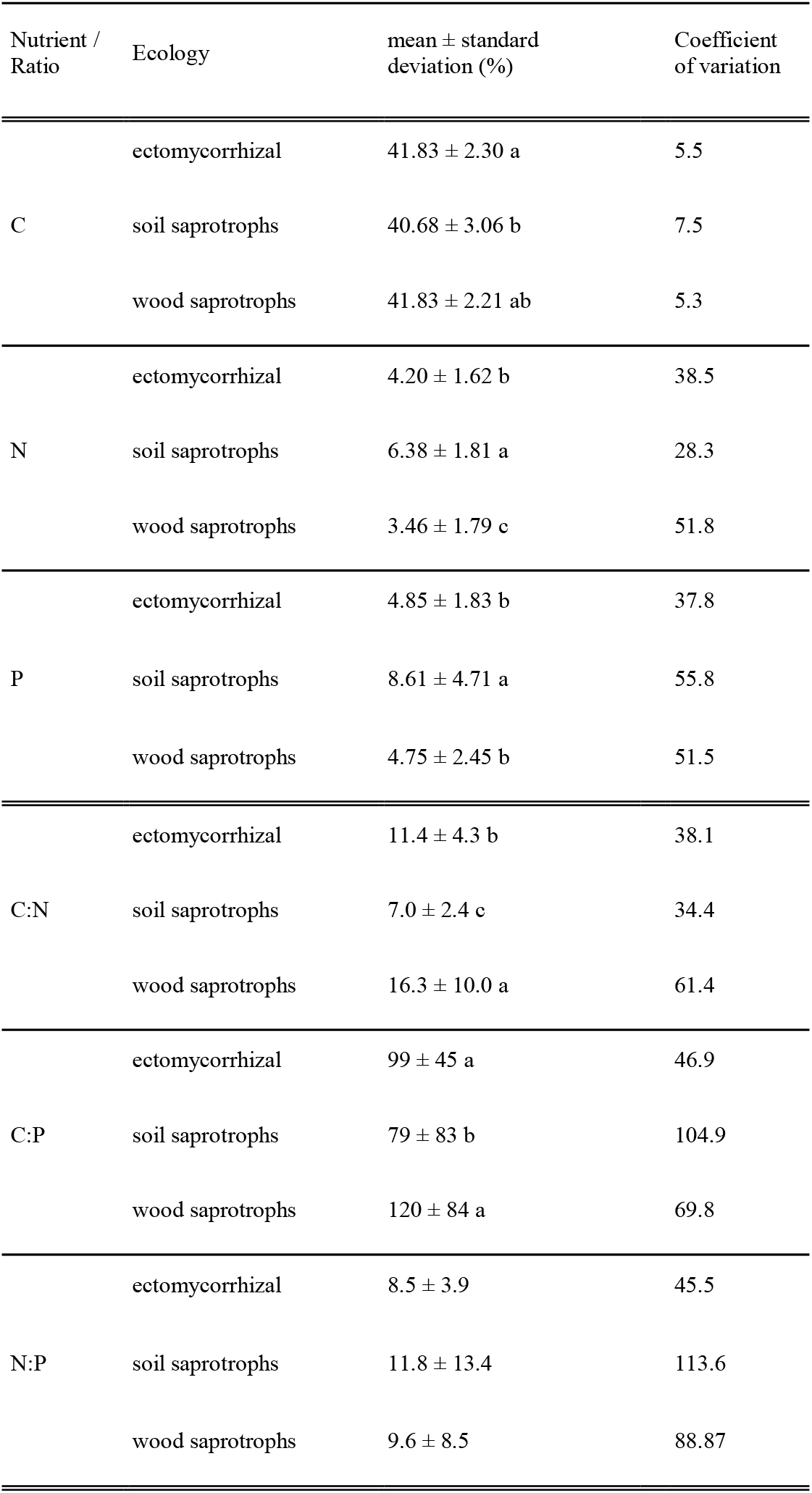
Nutrient content in ectomycorrhizal fungi, soil saprotrophs, and wood saprotrophs. Values labeled with distinct letters indicate significant differences (p < 0.05) among groups.

**Figure 3.**
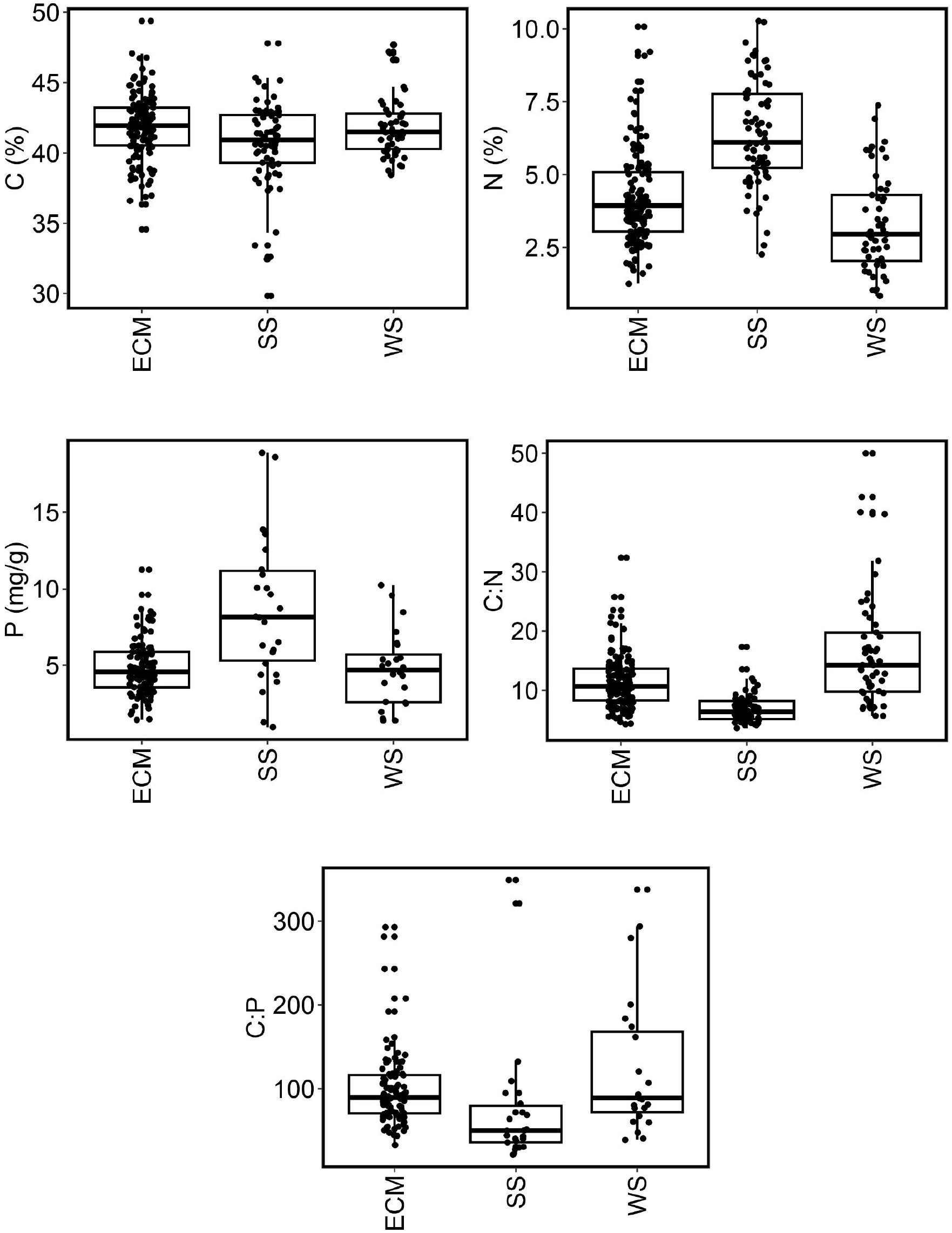
Nutrient content and nutrient stoichiometry in ectomycorrhizal fungi, soil saprotrophic fungi, and wood saprotrophic fungi. Boxplots indicate averages, lower and upper quartiles, and ranges (outliers excluded). ECM – ectomycorrhizal fungi, SS – soil saprotrophs, WS – wood saprotrophs.

### Phylogenetic conservation of nutrient content

The variation in nutrient content, as well as stoichiometric ratios of nutrients, increased from low taxonomic ranks to higher taxonomic ranks. While the coefficient of variation for C was 1.4 times higher at the order level compared to the species level, the corresponding values for N and P were approximately double and triple, respectively (Table 2).

**Table 2.**
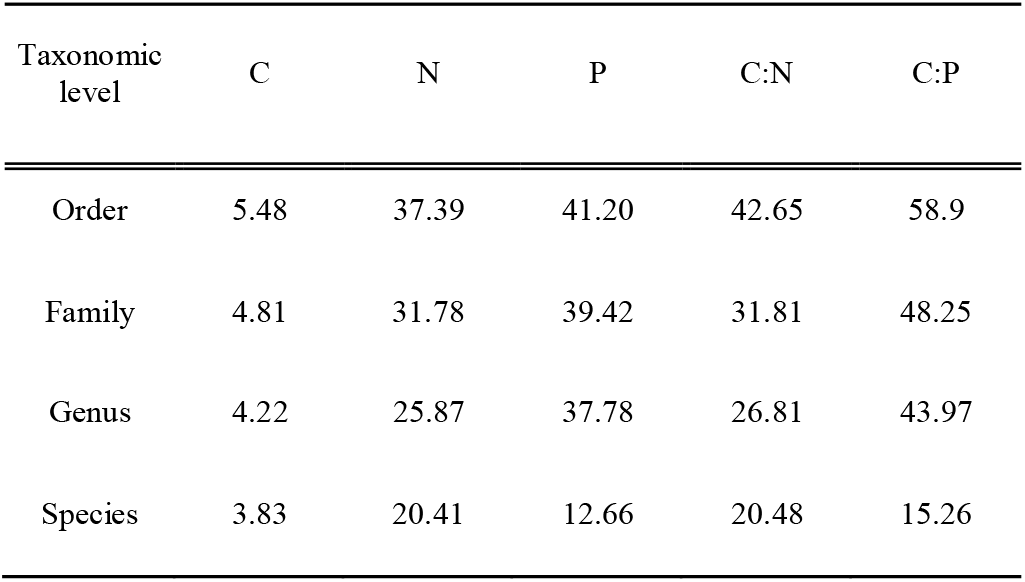
Variation of nutrient content and nutrient stoichiometry within taxonomic ranks of fungi, expressed as the mean coefficient of variation.

When analyzing the phylogenetic conservation of nutrient content traits in all samples, we found a significant phylogenetic signal for N and P content, as well as for the C:N and C:P ratios; no phylogenetic signal was found for C content (Supplementary Table 3). When testing lower taxonomic levels (order, genus), λ_P_ values differed between various taxa and generally decreased at lower taxonomy levels (although λ_P_ values were slightly affected by sampling depth, p = 0.08). The fact that phylogenetic conservation was observed for selected genera indicates that nutrient contents are phylogenetically conserved at the species level. This corroborates the observations of the limited variation of N and P content within fungal species and genera (Table 2).

The three most abundantly recorded orders – *Agaricales, Boletales*, and *Russulales* (138, 42, and 50 samples, respectively) – showed differences in the level of phylogenetic conservation of traits. While no phylogenetic signal was recorded in the *Russulales*, high levels of phylogenetic conservation for all three nutrients were observed in the *Boletales*.

## Discussion

Our results support the hypothesis that N and P content in fungal biomass differs among ecological guilds (Põlme et al., 2020; Root, 1967) – mycorrhizal, soil, and wood saprotrophic fungi. Thus, the nutrient content is, to some extent, determined by the environments to which these fungal groups have been adapted and the source of C that they use. In comparison to both saprotrophic groups, mycorrhizal fungi have an advantage of a stable supply of C as a result of their mutualistic relations with host plants (Bödeker et al., 2016). As a consequence of the trade with the host plants for C, N, and P, these fungi should pass the excess of N and P acquired from the soil to their host, retaining limited amounts of these nutrients in their mycelia (Kranabetter et al., 2019; Trocha et al., 2016; Zhang and Elser, 2017). This is reflected by lower contents of N and P in this guild, as well as lower variation in the content of these two nutrients. Conversely, the high average values discovered among soil saprotrophic species, where such nutrient trade does not take place, could be explained as an ability of soil saprotrophic fungi to store N and P in case of their excess (Kranabetter et al., 2019; Mooshammer et al., 2014; Trocha et al., 2016; Kalač, 2019; Quinché, 1997). The low content of N and P in wood saprotrophs compared to soil saprotrophs reflects their adaptation to the low nutrient contents in deadwood (Piché-Choquette et al., 2023; Watkinson et al., 2006).

Our results show – in addition to the effect of fungal lifestyle – the phylogenetic conservation of nutrient content. Lilleskov et al. (2011) showed that the response of fungi to elevated N content in soil is consistent only at the level of genera, and other ecological traits of fungi are also only rarely conserved at higher taxonomic levels (Zanne et al., 2020). This was also confirmed by the results of our study, where the conservation of nutrient content in biomass was highest at low taxonomic levels, while its variability increased with taxonomic rank. This may have been due to the fact that fungi have repeatedly evolved different lifestyles during evolution (Martin et al., 2016; Zanne et al., 2020), and therefore phylogenetic relationships at higher taxonomic levels may not correspond to the ecological features of fungal species. Although the differences in nutrient content between taxonomic groups are not as pronounced as differences between ecological guilds, phylogenetic relations seem to play some role in the determination of nutrient content in biomass.

Neither the phylogenetic relationship nor the membership in ecological guilds strongly determines the nutrient content in biomass, as indicated by the share of unexplained variation. This unexplained variation is likely associated with local, highly variable environmental conditions, where nutrient levels in fungi somewhat mirror those in the surroundings. This variability may also suggest an adaptive response to fluctuating environmental conditions (Camenzind et al., 2020; Vogt et al., 1981).

Due to dissimilar sources of N and P, their soil content could be highly variable. Such variability is associated with the type of geological bedrock, soil type, altitude, soil depth, and other factors; there can be many-fold differences in N content between various forest soils (Batjes, 1996; Huntington et al., 1988). Even higher is the variability of soil P content, exceeding differences of two orders of magnitude between various sites (Deng et al., 2017). Substantially low is nutrient content in wood, but the differences among tree species are considerable (Cowling and Merrill, 1966). Despite these dissimilarities and high, mutually independent variabilities of N and P causing considerable variability of their ratios within fungal habitats, the content of N and P in fungal biomass shows a significant positive correlation (Pearson’s r = 0.478, p < 0.01, Figure 2). No significant differences were found between all three ecological guilds in the N:P ratio, which could indicate that fungi in general maintain an approximately similar ratio of these nutrients, although the variability of this ratio is high, especially within both groups of saprotrophs. The N:P ratio is important particularly for mycorrhizal fungi, because it is tightly associated with their successful C, N, and P trade-off with the host plant (Johnson, 2010). This could explain the lower variability of the N:P ratio in the EcM group compared to the other two fungal groups (Table 1). Conversely, saprotrophic fungi may store both N and P independently of each other, according to their obtainable quantity, which could explain their higher variability in this ratio.

The content of chitin in fungal biomass was significantly lower in wood saprotrophic fungi than in the other two groups, although the variability in all three groups was high. As we hypothesized, the chitin content showed a positive correlation with the total N in mycelium (Pearson’s r = 0.298, p = 0.0239; Figure 2), indicating that fungi with sufficient N supply may synthesize more chitin. Chitin, as a chemical compound, contains approximately 6.9% of N (No and Meyers, 1995; Tracey, 1955), and N in chitin constitutes only 4.6-7.4% of the total N in the biomass (Supplementary Table 2).

Therefore, the contribution of N bound in chitin to the differences in N content among fungal ecological guilds is limited. These differences are more likely caused by N in proteins and intracellular content rather than by N localized in chitin. This is further supported by the observation that soil saprotrophic fungi, which have the highest levels of total nitrogen, do not necessarily contain the highest levels of chitin.

Fungal mycelium is an important component of food chains, serving as a source of nutrients after its decomposition. Differences in N content in biomass between fungi of all three ecological guilds are significant, indicating variable attractiveness of dead mycelium for subsequent decomposition. Not surprisingly, N content in fungal biomass was the best predictor of its decomposition rate (Brabcová et al., 2018; Fernandez et al., 2016). Due to higher N content, the biomass of soil saprotrophs should be more rapidly decomposed than that of the EcM and wood saprotrophic fungi.

The increasing deposition of N, which was frequently documented (Baldrian et al., 2023; Hůnová et al., 2017; Morrison et al., 2016), raises the question of its influence on fungi of various ecological guilds. While the wood, as the habitat of wood saprotrophic fungi, should be relatively unaffected by N deposition, the impact on soil saprotrophic and mycorrhizal fungi could be more pronounced.

Since the provision of N to plants is one of the essential functions of mycorrhizal symbioses (Kranabetter et al., 2019), increased N availability as a result of N deposition might affect the importance of mycorrhiza for plants. In such a case, at least the shift of the limitation to P could be expected. For the saprotrophic fungi, the direct impact of higher N availability should not be of cardinal significance since these fungi seem to be able to cope with various levels of nutrients and their ratios. But some indirect influence, such as increasing competition of other groups of microorganisms, could be assumed instead.

Our results indicate that both ecology and evolution determine nutrient contents in fungal biomass. In the light of ongoing global climatic change and continuing increased N deposition, it is especially important to assess how the differences in nutrient content among fungal guilds and the share of C that they receive within the soil affect the further fate of fungal biomass and whether it can impact organic matter accumulation or C storage in soils.

## Acknowledgements

This work was supported by the Czech Science Foundation (22-30769S) and by the Ministry of Education, Youth and Sports of the Czech Republic (CZ.02.01.01/00/22_008/0004635 - AdAgriF - Advanced methods of greenhouse gas emission reduction and sequestration in agriculture and forest landscape for climate change mitigation). The work of J.B. was supported by the Long-term Development Projects RVO67985831 and RVO61389005.

## Notes

### Competing Interest Statement

The authors have declared no competing interest.

